# Inferring functions of coding and non-coding genes using epigenomic patterns and deciphering the effect of combinatorics of transcription factors binding at promoters

**DOI:** 10.1101/2022.04.17.488570

**Authors:** Omkar Chandra, Madhu Sharma, Neetesh Pandey, Indra Prakash Jha, Shreya Mishra, Say Li Kong, Vibhor Kumar

## Abstract

The number of annotated genes in the human genome has increased tremendously, and understanding their biological role is challenging through experimental methods alone. There is a need for a computational approach to infer the function of genes, particularly for non-coding RNAs, with reliable explainability. We have utilized genomic features that are present across both coding and non-coding genes like transcription factor (TF) binding pattern, histone modifications, and DNase hypersensitivity profiles to predict ontology-based functions of genes. Our approach for gene function prediction (GFPred) made reliable predictions (>90% balanced accuracy) for 486 gene-sets. Further analysis revealed that predictability using only TF-binding patterns at promoters is also high, and it paved the way for studying the effect of their combinatorics. The predicted associations between functions and genes were validated for their reliability using PubMed abstract mining. Clustering functions based on shared top predictive TFs revealed many latent groups of gene-sets involved in common major biological processes. Available CRISPR screens also supported the inferred association of genes with the major biological processes of latent groups of gene-sets. For the explainability of our approach, we also made more insights into the effect of combinatorics of TF binding (especially TF-pairs) on association with biological functions.

## Introduction

A biochemical pathway in a cell includes the role of both coding and non-coding RNA (ncRNA) genes’ products. The functional role of non-coding RNA is prominently in trans or cis-regulation of the coding genes whose products are the backbone of biochemical pathways (Rinn and Chang 2012). The ncRNA genes are short (around 200 base pairs long) and share homology across many genes, and they have various molecular mechanisms through which they exert their functions in a myriad of molecular and cellular functions; because of these reasons, it is hard to study ncRNAs experimentally (Kevin C. Wang 2011; X. Zhang et al. 2019). A promising way to dissect the functions of the ncRNA genes is through computational analysis by leveraging the existing knowledge of gene-function or gene-disease relationships. Gene ontologies represent empirically annotated relationships between disease, functions, and genes. In the past, multiple research groups have utilized these ontologies for predicting associations of genes with functions and diseases(Zhao et al. 2020). Here we have used the word “function” to represent gene-sets of molecular functions and biological processes for ease of reading.

Predictive models are good at identifying similarities between data points and are extensively used in gene function identification by comparing the features of unknown and known genes. However, using the most relevant biological signals as features for training a predictive model is still a crucial factor. The features of the known genes must essentially represent the function well for robust prediction. A straightforward approach to gene function prediction is by comparing the primary nucleotide sequences of genes and proteins of known function with the genes of unknown function (Zhao et al. 2020; Hanyu Zhang et al. 2019; Kulmanov and Hoehndorf 2019; N. Zhou et al. 2019), but it has been shown that alternative isoforms can be functionally divergent (X. Yang et al. 2016) and primary sequence comparison for non-coding genes would be of limited use because of the lack of reference to non-coding genes whose biological functions are known. Some researchers have used the ontological relationships of genes to identify disease-related ncRNA genes (P. Yang et al. 2012), but the number of annotations of ncRNA genes in the ontologies is less and would result in less coverage. Few studies have used gene expression data as features for identifying the non-coding genes based on the co-expression of the coding genes (Liu et al. 2018). However, a vast number of functionally unrelated genes can show correlation at a given instant, and genes involved in the same pathway may not exhibit any correlation (Uygun et al. 2016). Thus, current approaches fail to effectively utilize the right input genomic features to predict non-coding gene functions.

It is well known that non-coding RNAs regulate the transcription of genes involved in the same biological process through interactions with chromatin, RNA, and protein (Sun and Wong 2016). However, the question is how to predict their association with any function. Genomic features like the binding of transcription factors (TFs) are present across both coding and non-coding genes as they are required to modulate the gene expression (Venters and Pugh 2013; Yan et al. 2021). Epigenetic marks and chromatin structure work in tandem with the TFs in the modulation of gene expression (B. Li, Carey, and Workman 2007). In the past, epigenome profiles have been used to predict gene expression (Kumar et al. 2013) and the association between disease and single nucleotide polymorphism (SNP) (Tak and Farnham 2015). At the same time, many previous studies which published TF ChIP-seq profiles have tried to associate binding patterns of 1 or 2 TFs, at a time, with genes of a particular function. Using TFs as features can also help make insights into the combinatorics (synergy and cooperativity) involved in regulating different functions (Venkatesh et al. 2021). However, a comprehensive analysis of combinatorics of binding patterns of large numbers of TFs at promoters and their associations with the function of genes has rarely been done.

Here, we devised an approach to use combinatorics of epigenomic signals, especially TF binding patterns at promoters of genes to predict the ontology-based function of genes. Accordingly, to capture all the signatures of factors involved during the modulation events that would occur during the transcription of a gene; we leveraged a large number of publicly available ChIP-seq data of TFs, histone modification marks, and DNase I hypersensitivity sites along with cap analysis gene expression (CAGE) tags to include the expression of genes including non-polyadenylated ones. In order to gain more insight into the reliability of our method, we performed downstream analysis involving top predictive ChIP-seq profiles for clustering of functions and associating genes with those clusters of gene-sets. We also made insights into the specificity of simple combinatorics of TFs (i.e TF-pair) towards functions.

## Results

We developed our approach based on the hypothesis that the coordinated expression and functional association of genes are brought about by a few common key regulatory factors present across both coding and non-coding genes. We downloaded ChIP-seq profiles of TFs, histone modification marks, and DNase I-seq and CAGE-tags from different sources and estimated their read-count within 1 Kb of transcription start sites (TSS) of genes, in other words, 2Kbp wide region around promoter. The flowchart of our approach (GFpred) is shown in Figure 1.

**Figure 1:**
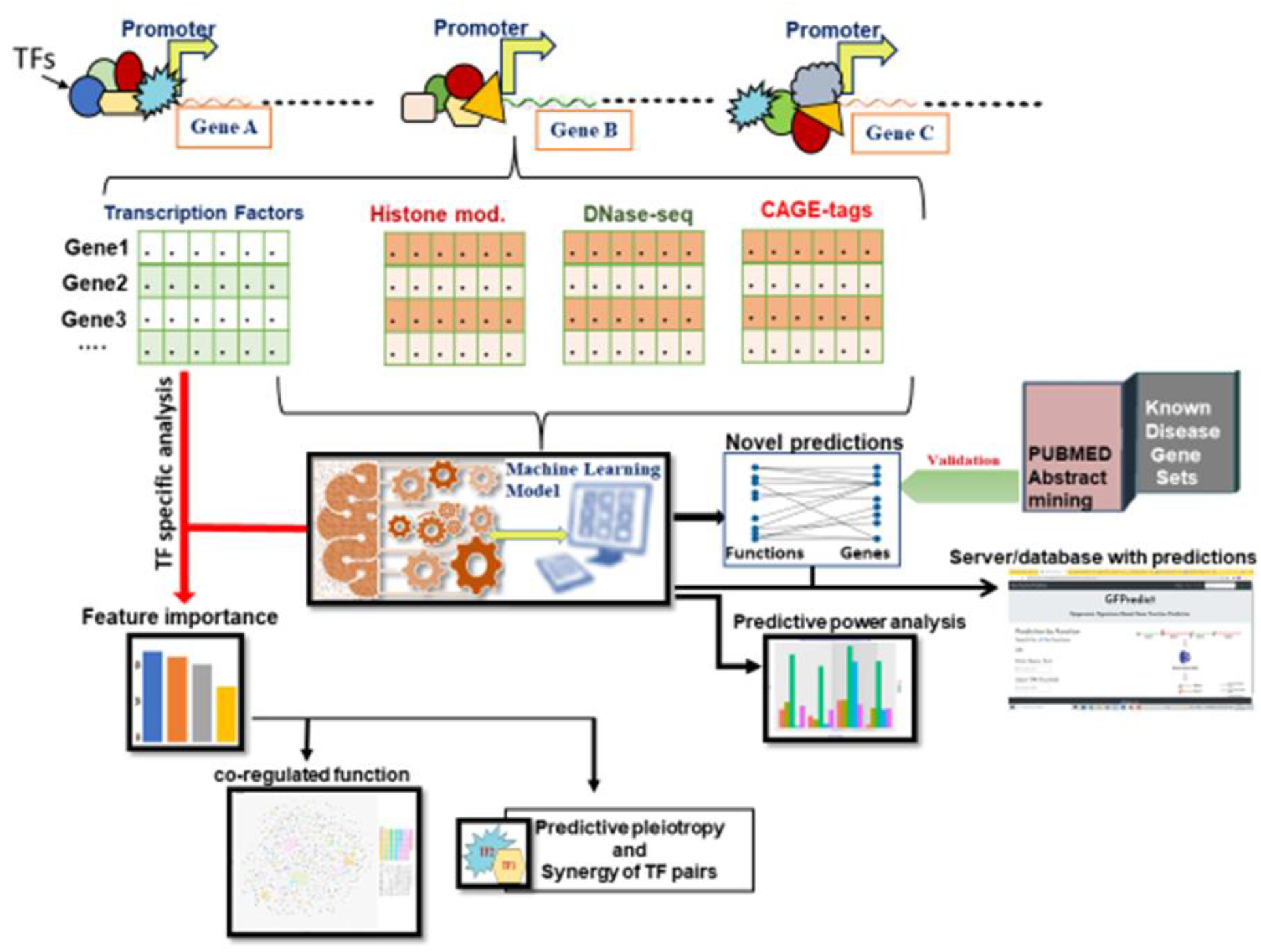
Flowchart of our analysis to predict gene functions using epigenome profiles.

### Gene functions are predictable using the epigenomic signals at the promoter regions

Machine learning (ML) algorithms were trained for each of the biological functions of the ontologies. We trained 5 different ML models using TFs binding patterns and other features (ChIP-seq data of DNase hypersensitivity regions, histone marks, CAGE tags) for a total of 9559 function gene-sets downloaded from the MSigDB database (Liberzon et al. 2011). We performed two approaches for predictive modelling. We used 823 TF ChIP-seq libraries from normal (non-diseased) samples for estimating feature scores in the first approach. For the other approach, we used ChIP-seq profiles of TFs and histone modifications ChIP-seq (n = 621) and DNase-seq (n = 255) and CAGE tags (n = 255) from non-diseased samples. While using the second approach, we achieved good predictions for many functions, such that using randomForest; the sensitivity was above 80% and minimum specificity of 90% for 425 genesets. Other four ML models (linear regression, logistic regression, SVM, and XGBoost) showed 100 to 300 gene-sets with a sensitivity of 80% and specificity of 90% (Figure 2A). However, when using only 823 TF-ChIP-seq profiles, the number of functions with similar predictability did not reduce substantially. Using the threshold criteria of 80% sensitivity and 90% specificity, we had 318 functions using randomForest. We took the union of functions with good predictability (sensitivity > 80 %, specificity > 90%) from 5 ML models. We found that using only TF ChIP-seq, 670 functions had good predictability from at least one of the five ML models. However, using all features (TF, Histone modification, CAGE-tags, DNase-seq) we had an increase of only 15% in the number of functions (total number = 773) (Figure 2B) with good predictability (sensitivity > 80%, specificity > 90%) with at least one ML model. When we used the criteria of sensitivity greater than 70%(with specificity > 90%), the number of functions based on union from 5 ML models was above 1300 with TF ChIP-seq as features (Supplementary Figure 1). The balanced accuracy for prediction for a few functions is shown in Figure 2C. Evaluation metrics used for each gene-sets can be checked in the Supporting File 1.

**Figure 2:**
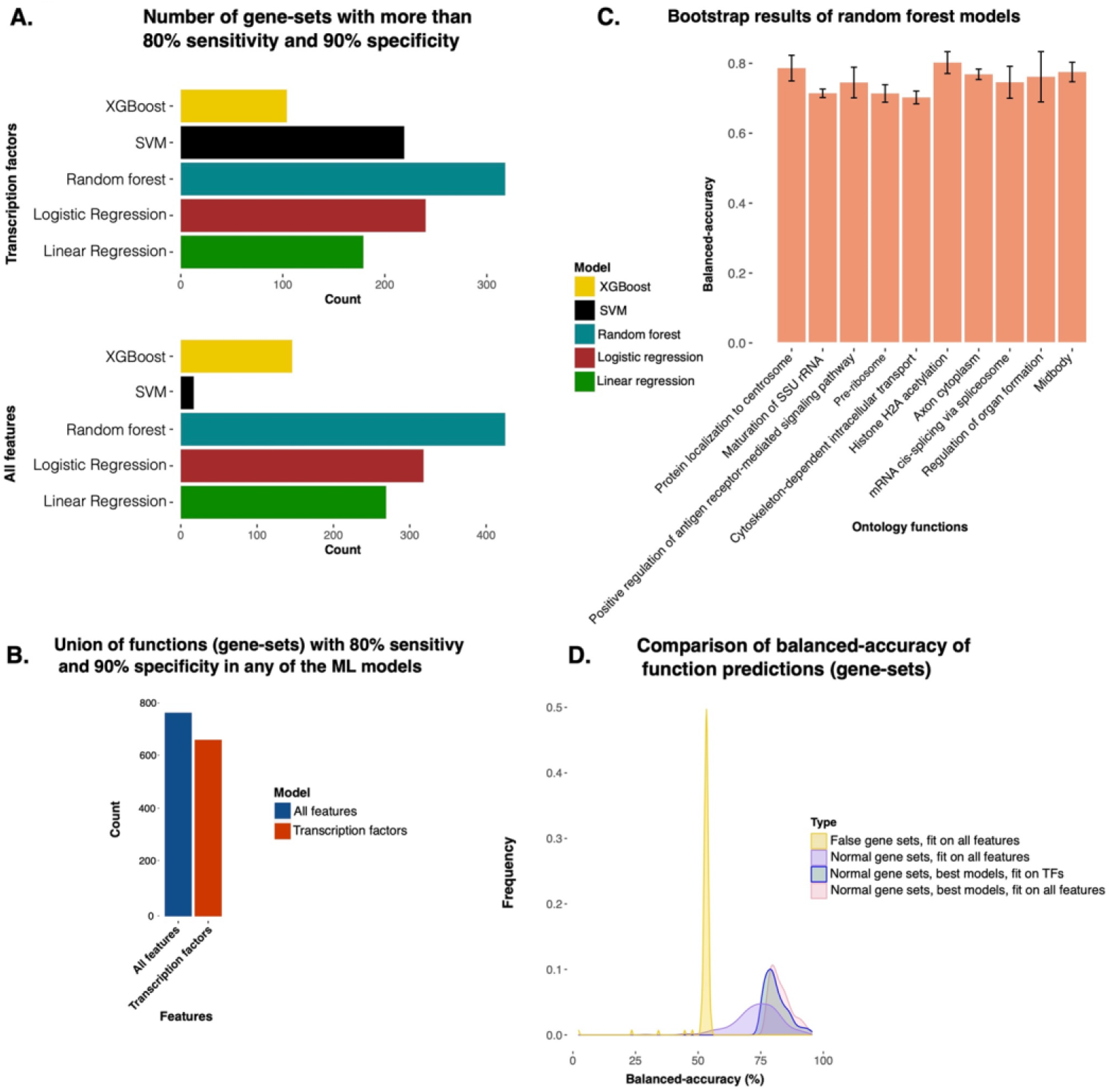
An overview of the predictive power of epigenome profiles, especially transcription factor binding patterns at promoters for predicting gene function. A) Bar plot showing the number of functional gene-sets which had good predictions on the test set (80% sensitivity and 90% specificity) using 5 different machine learning (ML) models. The upper panel shows the number of functions with the good prediction by ML models using 853 transcription factor (TF) ChIP-seq profiles. The lower panel shows the ML model using 5 different types of profiles (TF and histone modification ChIP-seq, DNase-seq, CAGE-tags). B) The bar-plot shows the number of union set of functions with good predictability (80% sensitivity and 90% specificity) using any of the 5 ML models. C) The balanced accuracy was achieved for a few functions using bootstrapping (iterations = 5). D) A plot to show the sanity of our approach. Here the density plot in yellow color shows the distribution of balanced accuracy achieved with false gene-sets (gene-sets created by random sampling). Other density plots show the distribution of balanced accuracy achieved using empirically annotated gene-sets. The density plot for some functions with balanced accuracy above the 35 percentile among all the functions is also shown.

#### Non-random nature and relevance of high predictability

To ensure that high predictability achieved using our approach is non-stochastic, we constructed a null model as a control. For this purpose, we checked if modeling is possible on ‘false gene-sets,’ apart from the gene-sets annotated empirically. We created 200 false genesets by randomly shuffling the genes from existing gene-sets. The best performing random forest algorithm trained on these false gene-sets showed an overall balanced accuracy of not more than 55% on average. In comparison, the balanced accuracy of the models on the empirically annotated gene-sets is 75% on average, as shown in Figure 2D. Our result indicates that good predictability is possible only for biologically relevant gene-sets, and there is an inherent pattern of regulation exhibited by a few common regulators at the promoter sites of the genes associated with the same biological function, supporting our hypothesis.

### Inference from clustering of functions

There are few indirect studies on the coregulation of functions by the combinatorics of TF binding (Wu and Lai 2016). However, we used a direct approach of studying the effect on function predictability due to combinatorics of TFs to get an insight into major functional groups of coding and non-coding RNA. We found 50 (Figure 3A) prominent clusters of functions (Supporting File 2) based on shared top predictors. In addition, we found that in some clusters, the majority of the functions were involved in similar major biological activity (Supporting File 2). For example, one of the large clusters (cluster-47) related to the cell cycle includes ‘regulation of cell cycle process’, ‘cytokinesis’, ‘microtubule-organizing centre’, ‘nucleolus’, ‘regulation of cellular protein localization’ (Figure 3A). Some of the top predictive transcription factors and cofactors shared among the functions of cluster-47 are *CTCF, XRN2, BRD4, SMARCA4, and PARP1* (Figure 3B). The role of *CTCF, MYC, PARP-1*, and *SMARCA4* in cell cycle regulation has been reported by previous studies (Hyle et al. 2019; Haoyue Zhang et al. 2021; L. Yang et al. 2013; Hendricks, Shanahan, and Lees 2004). One other cluster (cluster-26) shown in Figure 4A consisted of early development and morphogenesis-related terms. Some of the shared top predictors for functions in cluster-26 included POU5F1 (Bakhmet and Tomilin 2021), RNF2, and SMARCB1 (Meurer et al. 2021; Kenny et al. 2021), SIX1 (Meurer et al. 2021), which are known to regulate genes involved in early development. The results show that cluster members (genesets) have unrelated functional roles but a non-discernible role in the overall major biological activity. For example, in cluster-47, the majority of the members of the clusters have an apparent role in the major biological function–the cell cycle and ‘negative regulation of catabolic process’, is one of the members of the cluster with a generic cellular function but the fact it is one the members sharing few of the regulatory elements with other members imply its indirect role in the cell cycle. Such indirect role of functions in major biological processes can be deduced in other clusters (Supporting File 2). Thus the emergence of clusters of functions broadens the scope of linking genes to major biological processes and hints at the specificity of the binding patterns of the regulators (TFs and cofactors).

**Figure 3:**
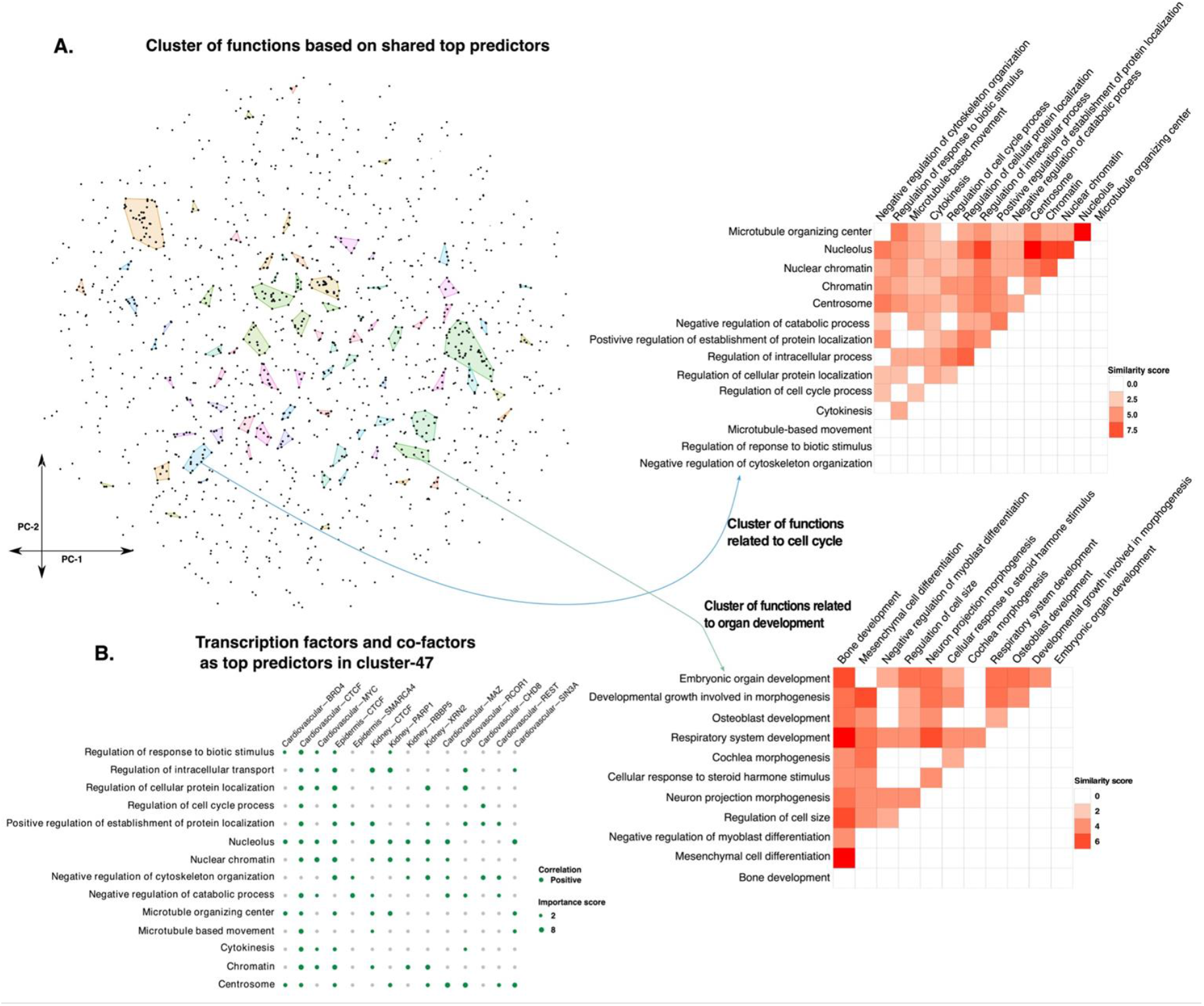
Clustering of functions based on shared predictive epigenomic features reveal their potential overlap of involvement in major cellular processes. A) tSNE plot and visualization of DBSCAN-based clustering of function (gene-sets). Here, every dot in the tSNE plot shows a gene-set. The details about the two clusters are displayed as a heatmap showing the similarity in terms of the number of common top predictors (ChIP-seq profiles in top 20 predictors). It can be noticed that most of the functions in one of the clusters are involved in the cell-cycle, such as B) The dot plot shows the value of feature importance of ChIP-seq profiles of TFs and cofactors for functions belonging to one of the clusters (cluster-47). The feature importance value not lying in the top 20 is shown as minimum dot size.

**Figure 4:**
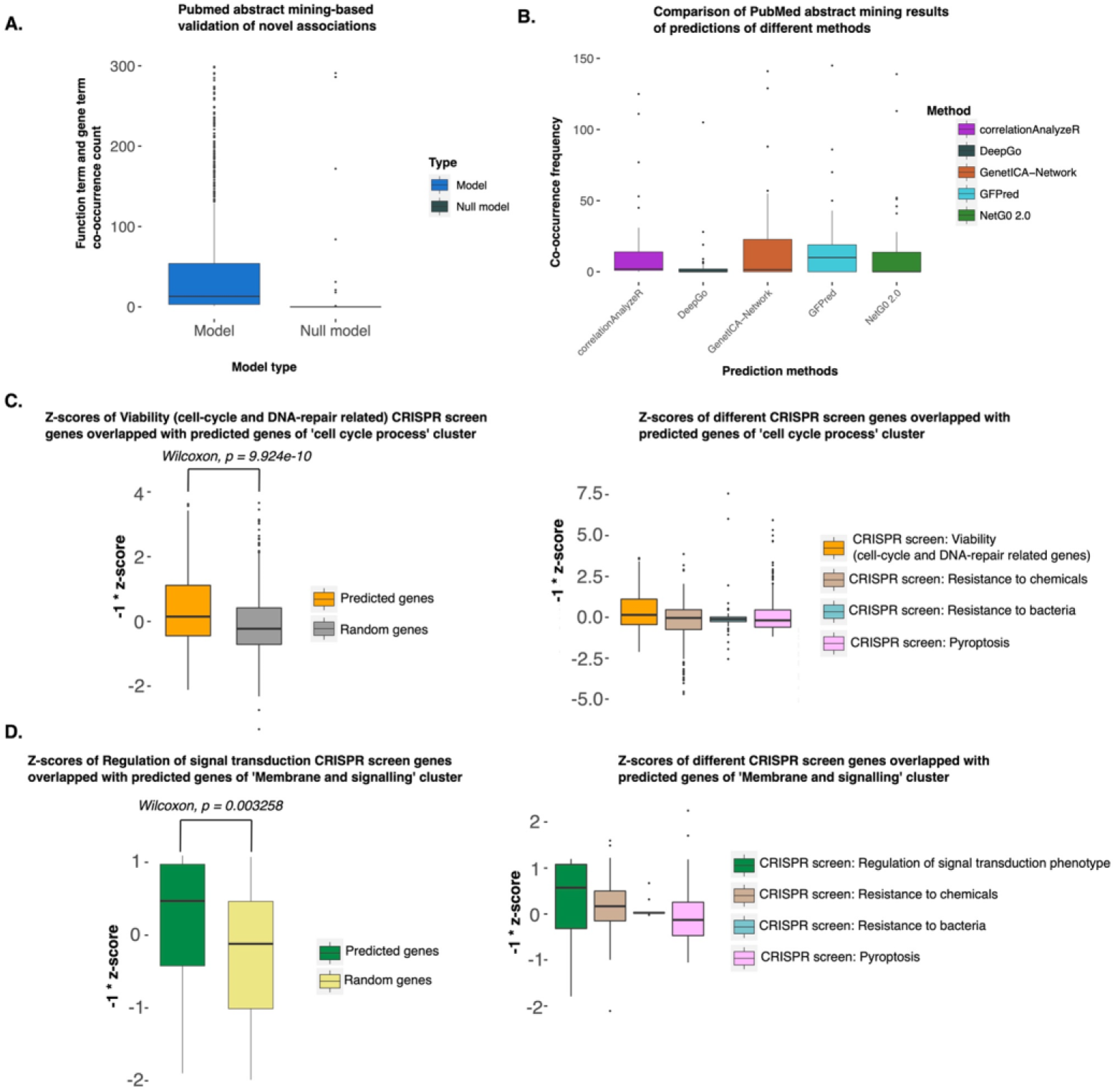
Validation of predictions of novel association between function and genes. A) The box-plot shows the frequency of co-occurrence of function terms and gene names in PubMed abstracts. The left box plot shows the frequency of the novel predictions made by GFPred, while the right one shows random pairs of functions and genes. The novel and random associations between function and genes were not present in the gene-sets we used for training or testing. B) Benchmarking and comparing function prediction methods for finding an association between function and gene. C) Validation using CRISPR screen: ‘Viability’ for identifying genes involved in cell-cycle and proliferation D) Validation using CRISPR screen: ‘Regulation of signal transduction phenotype’ to identify genes involved in immune response signalling pathways.

### Independent validations of the predicted results

#### Pubmed abstract mining of co-occurrence of gene term and function term

To check if the predicted results are of any biological relevance, the co-occurrence of the predicted gene term and the corresponding biological function term of the ontology is searched in the abstracts of the PubMed articles published from 1990 to 2021. The boxplot in Figure 4A shows the total co-occurrence of predicted gene term and function term pairs compared against random gene term and random function term pairs as control. This result adds to the confidence in our predicted results.

#### Related disease-gene links

We also checked if the predicted genes are reported in related human disease gene sets. We found 23 intersections between the predicted genes and the disease gene-sets obtained from the DisGeNET database (Piñero et al. 2017) (Supplementary Table 1). Refer to Supplementary Methods for the procedure followed to perform the intersection. There were a total of 15 intersections with the Mouse Genome Informatics (MGI) (Blake et al. 2021) (Supplementary Table 2).

### Comparison of predicted results with other gene-function prediction methods

Gene function prediction is one of the classical problems in computational biology. Some of the recent methods to predict the ontology-based functions of genes have utilized different features like primary amino acid sequence (NetGo 2.0, DeepGo), gene expression (correlation AnalyzeR), and network inference using co-functionality of genes obtained through transcriptomic profiles (GenetICA-Network) (Miller and Bishop 2021; Kulmanov et al. 2018; Urzúa-Traslaviña et al. 2021; Yao et al. 2021). We compared the abstract mining results on the predictions of these methods against the novel associations inferred by our approach (Figure 4B). The co-occurrence of input ontology term and predicted gene term at least once in the PubMed abstracts for the 50 gene-sets of our method is significantly more compared to the same input gene terms and their predicted ontology terms by AnalyzeR, DeepGo, NetGo 2.0, and comparable to GenetICA-Network. However, the median of the frequency of co-occurrence of input terms and corresponding related terms predicted by our method is more than all the other methods. Note that we compared the prediction results and not the performance of the models of different gene function prediction methods because our implementation’s input features and approach for gene function prediction are completely different from the other methods. Therefore, benchmarking is possible only on the predicted novel associations with functions for a group of genes.

### CRISPR-based validation of association of genes with major biological processes of clusters of functions

Our approach of grouping functions based on common top predictors (TFs or cofactors) leads to new way of finding links (direct and indirect associations) between coding and non-coding genes with a few major biological processes. In order to evaluate the results of the discovery of such new links between genes and major biological processes, we analyzed available CRISPR screens. First, we used the CRISPR screen: ‘viability’ in human pluripotent stem cells (hPSC), where the hPSC-enriched essential genes appeared to be mainly encoding transcription factors and proteins related to cell-cycle and DNA-repair (Yilmaz et al. 2018). The novel predicted genes in functions belonging to cluster-47 (mainly associated with cell-cycle processes) had significantly higher z-scores in comparison to an equal number of random genes in the same CRISPR screen for viability of hPSC (Figure 4C). However, the novel predicted genes for cluster-47 had comparatively less z-scores in other CRISPR screens: ‘resistance to chemicals’, ‘resistance to bacteria’ and ‘pyroptosis’ (Schinzel et al. 2019; Jeng et al. 2019; Alimov et al. 2019). For another validation, we used the CRISPR screen: ‘regulation of signal transduction phenotype’, which highlighted critical components of the tumor-immune synapse and the importance of cancer cell interferon-γ signaling in modulating NK activity (Pech et al. 2019). A cluster (cluster-52) consisted of functions related to immune cell activation and its associated pathways (see Supporting File2 for other members) during clustering based on top predictors. The novel predicted genes of cluster-52 had higher z-scores compared to an equal number of random genes in the same screen for regulation of signal transduction phenotype. However, the novel predicted genes for cluster-52 had comparatively less z-scores in other CRISPR screens (Figure 4D). CRISPR screens’ validations assert the associations of novel genes with major biological processes and link the underlying regulatory factors (top predictors) to those biological processes.

#### Insight about the effect of binding patterns of TF pairs for better inference of functions

Our prediction approach using multiple TF ChIP-seq read-count at promoters of genes seems reliable; however, there is still a need to study combinatorics of TF binding for better explainability. Hence, to gain more explainability and reliability in our approach, we tried to understand the effect of the simplest combinatorics of TF-pair binding patterns. Transcription factors have pleiotropic effects even within the same tissue or cell-type (Chesmore et al. 2016). As expected, a few TFs had high feature importance scores for many functions. To analyze the predictive pleiotropy of TFs, we searched for TF ChIP-seq pairs (irrespective of cell-types), which emerged together as top predictors of different functions. A few TF ChIP-seq pairs were among the top predictors of multiple functions (Figure 5A and Figure 5B). The occurrence of a TF pair among the top important features across multiple biological functions indicated their pleiotropic predictive power (Wang, Liao, and Zhang 2010). Further, we checked for the diversity of functions for which they were top predictors for every TF pair. For diversity estimation, we counted TF pairs occurrences in the clusters of co-regulated functions (Supporting File 3). We performed the same task 2 times; the first time, we used TF ChIP-seq pairs irrespective of the cell-type, and the second time, we used TF ChIP-seq pairs from the same cell-type. We found that TF-pairs appeared to have predictive pleiotropy for many functions but had less diversity in terms of clusters of co-regulated functions. ChIP-seq patterns of BATF and RUNX3 at promoters in B cells (GM12878) appeared together among the top 20 predictors for 11 functional gene-sets. However, these 11 gene-sets belonged to only 2 clusters of functions mainly involved in immune cell activation and differentiation (Supporting File 4). Similarly, DNA-binding profiles at promoters in adipocytes by CEBPA and E2F4 appeared to be top predictors for 8 gene-sets (functions) belonging to a single cluster and mainly associated with response to the stimulus by peptides (like insulin) and monosaccharides and related metabolic processes. Thus, some of the TF-pairs seem to have more specificity toward major cellular or biological processes that can be exploited to confirm the prediction of coding and non-coding genes.

**Figure 5:**
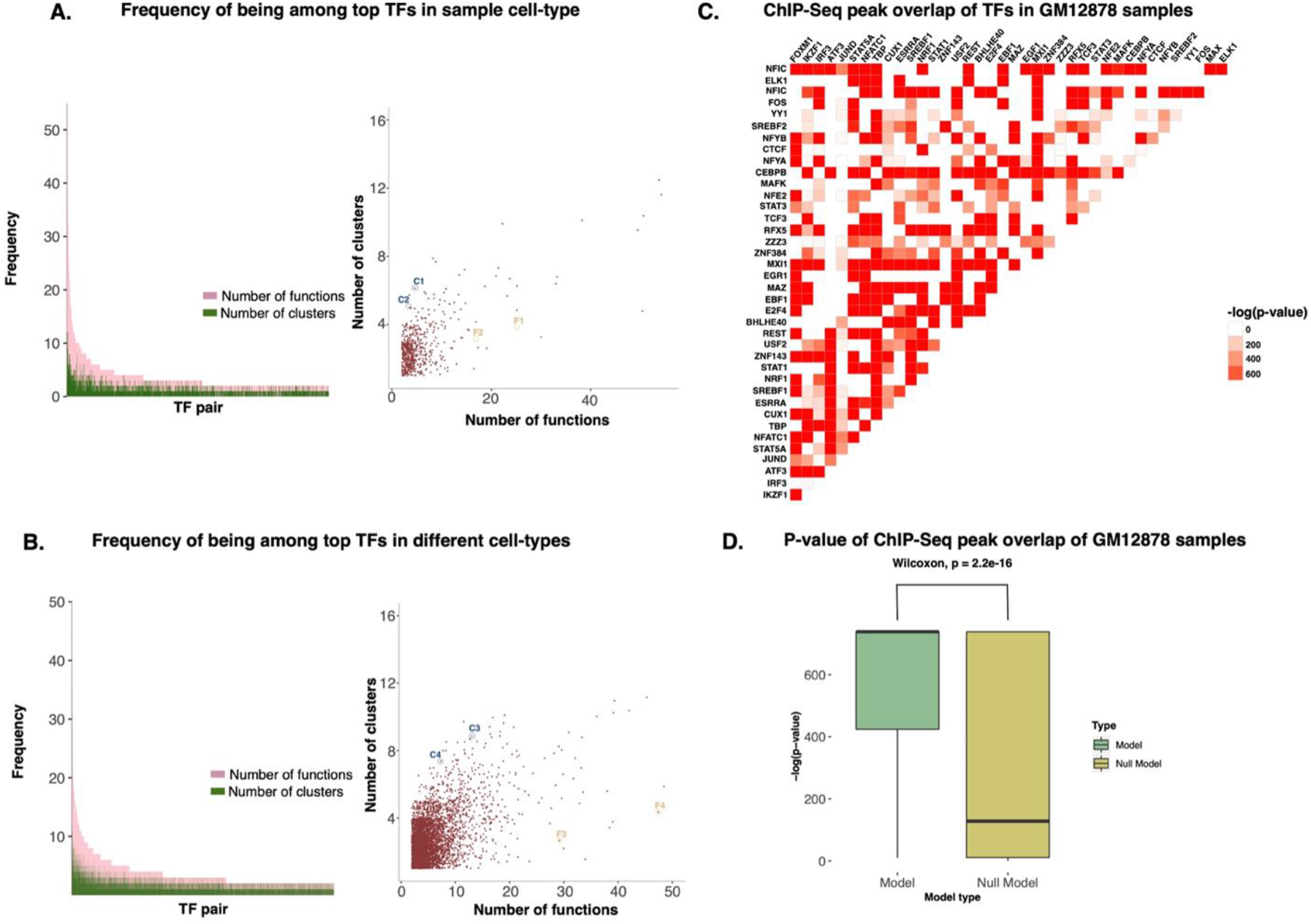
Insight into the co-occurrence of Transcription factor (TF) pairs among predictors and their synergy. A) The count of functions (pink) and the clusters of functions (green) for which TF ChIP-seq pairs appeared among the top 20 predictors. The panel on the right shows the same counts as a scatter plot. Here the TF ChIP-seq in a pair were allowed to be from different cells-type. The TF-pairs shown with symbols are C1: REST-GABPA, C2: ZNF76-TET2, F1: STAT1-IRF1, F2: LHX2-ZNF92. B) The count of functions and their clusters for which pairs of TF ChIP-seq profiles from the same cell-type appeared together as the top predictors. The panel on the right shows the scatter plot version of such counts. The TF-pairs shown with symbols are C3: E2F4-GATA1, C4: MAZ-GATA1, F3: ZNF366-SPI1, F4: SPI1-STAT1. C) Heatmap showing the significance of overlap of TF ChIP-seq peaks in GM12878 cells at promoters. D) The box-plot of values of significance (-log(P-value)) of overlap of promoter-peaks of TF ChIP-seq pairs in GM12878 cells which appeared together as top predictors in one or more functions. On the right is the box-plot of the significance of overlap (Wilcoxon rank-sum test, p-value < 2.2e-16) of promoter peaks for random pairs of TF ChIP-seq profiles in GM12878 cells.

Only a minor group of TF-pairs were top predictors of the gene-set belonging to more diverse clusters of co-regulated functions. Especially pairs involving CTCF showed more diversity in a cluster of co-regulated functions. The TFs, ZCAN5FB, and CTCF appeared as the top predictors for functions belonging to more than 12 co-regulated clusters. Similarly, TET3 and CTCF appeared as the top predictor of functions from 6 clusters. CTCF is known to have a more general effect than other TFs. CTCF is known to play the general role of insulator (Kim, Yu, and Kaang 2015). Nevertheless, its co-occurrence with certain TFs as the top predictor also highlights another possible role in various cellular processes.

In order to further make insight into the non-random aspect of co-occurrence of TF pairs (Figure 5C) as top predictors, we investigated the overlap of the peaks of their ChIP-seq profiles. It was based on the notion that if TF-pair occurrence as top predictors has no relation with conserved biological processes, then the overlap of their peaks would appear as a random event. For this purpose, we used the R package ‘*ChIPpeakAnno*’ (L. J. Zhu et al. 2010) and analyzed TF ChIP-seq peaks in the GM12878 cell line. We compared the overlap of co-predictor TF-pair ChIP-seq peaks in the same cell-type with random TF pairs as control. Here, co-predictor TF-pair were defined as, pair of TF ChIP-seq profiles in GM12878 cells, which appeared among the top 20 predictors (co-predictive) for any function (Supplementary Figure 2B). We found that the enrichment of overlap of peaks at promoters in GM12878 for such co-predictive TF pairs was much more significant than random TF ChIP-seq pairs (Figure 5D). Such observations build confidence in our approach and indicate that the top predictors’ analysis offers insights into TF-TF synergy through higher co-binding frequency at promoters of the genes involved in the same biological functions.

### Insights about functions of non-coding RNAs

From the results of ML models, we relied on predictions (with >60% confidence score) for 1,200 long non-coding genes, associating them with various biological processes and molecular functions. Table 1 and Supplementary Table 3 contain the list of non-coding RNAs with corresponding literature support. Our results for predicted major cellular processes for some non-coding RNA based on function clusters are provided in Supporting File 5.

**Table 1:** List of predicted functions of non-coding RNAs with experimental evidence.

| Ontology | Predicted non-coding gene | Literature evidence |
| --- | --- | --- |
| GO_STEROL_HOMEOSTASIS | LINC02356 | (Raulerson et al. 2019) |
| GO_HEART_DEVELOPMENT | AP001528/ENSG00000280339 | (Y.-X. Chen et al. 2021) |
| GO_EYE_DEVELOPMENT | AC078909.1 | (Donato et al. 2020) |
| GO_PHOSPHOLIPID_METABOLIC_PROCESS | ENSG00000257023 | (Elaine Hardman et al. 2019) |
| GO_SYNAPSE_ORGANIZATION | MIR4281 | (P. Zhu et al. 2022) |
| GO_NEURON_MATURATION | ENSG00000274367 | (Z. Li et al. 2021) |
| GO_NEGATIVE_REGULATION_OF_INTERLEUKIN_6_PROD<br>UCTION | LINC00528 | (Sage et al. 2020) |
| GO_REGULATION_OF_HOMEOSTATIC_PROCESS | MIR658 | (Sánchez-Jiménez et al. 2013) |
| GO_KERATINOCYTE_DIFFERENTIATION | PAUPAR | (J. Chen et al. 2020) |
| GO_IN_UTERO_EMBRYONIC_DEVELOPMENT | MIR5001 | (Whittington et al. 2018) |

#### Case-1: The role of lncRNA AC078909.1 in retinitis pigmentosa

Retinitis pigmentosa is one of the most heterogeneous inherited disorders. Luigi *et al*. (Donato et al. 2020) designed the experiment to understand the molecular mechanisms of oxidative stress underlying the etiopathogenesis of retinitis. Their RNA-Seq-based experiment using human retinal pigment epithelium cells treated by the oxidant agent N-retinylidene-N-retinylethanolamine revealed lncRNAs associated with various pathological conditions leading to retinal cell death. Out of two down-regulated lncRNAs with the highest fold change in experiments by Luigi *et al*., one of those genes (AC078909.1) was also predicted to have a role in eye development by our method.

#### Case-2: The role of LINC00528 as one of the mediators of lung tumor immune response

The tumor immune microenvironment is a crucial mediator of lung tumorigenesis. Adam *et al*. (Sage et al. 2020) sought to identify the landscape of tumor-infiltrating immune cells in the context of long non-coding RNA (lncRNAs). They analyzed lncRNA profiles of lung adenocarcinoma tumors by interrogating the single-cell RNA sequencing data from microdissected and non-microdissected tumor samples. The predicted lncRNA LINC00528 in ontology ‘negative regulation of interleukin-6 production’ is one of the top 10 lncRNA genes positively correlated with *PTPRC* protein, a marker for immune cells (in the study by Adam et al.). Such results indicate the potential role of LINC00528 in the infiltration of immune cells in immunogenic tumors.

## Discussion

To predict the function of non-coding RNAs, researchers would have to use new kinds of assays or genomic features in prediction systems. We used epigenomic and CAGE-tags profiles to predict their functions to have a common set of features between coding and non-coding genes. Using the union of different ML models, we achieved good predictability for more than 780 functions with all features and 650 functions using 853 TF and cofactors ChIP-seq. We independently validated our results using different datasets. We also compared our method to other methods that use various other features for function prediction. Our results hint that in the future, with an increase in the availability of ChIP-seq profiles, the number of functions with good predictability will also grow.

Further downstream analysis revealed an interesting pattern that the majority of the functions which shared the same top predictors (especially ChIP-seq profile from the same cell-type) were either related to similar major cellular processes or had some dependencies on each other. Thus, despite having a seemingly unrelated biological role, functional gene-sets showed convergence in terms of association with major cellular processes like cell cycle and transport. Such observation is because of shared similarity in patterns of some epigenomic (or TF-binding) features at promoters of genes. On the same logic, if a few epigenomic features appear to be important common determinants (or predictors) for two known gene-sets, the genes of those gene-sets could likely be involved in some major function. In order words, our analysis goes beyond the boundary of currently defined gene-sets of function to highlight the effect of TFs. For example, for cluster-47, the top predictors are MYC, PARP-1, CTCF, and SMARCA4, which are involved in cell-cycle (Hyle et al. 2019; Haoyue Zhang et al. 2021; L. Yang et al. 2013; Hendricks, Shanahan, and Lees 2004). Thus, our analysis of the co-prediction (coregulation) of functions through top predictors shows the interdependence between functional gene-sets and may explain the perturbation’s effect on a key regulator that can potentially affect a myriad of functions. Two major aspects highlight the novelty of our study; i) deciphering combinatorics of TF-binding at promoters for association with functions ii) grouping of known gene-sets using top co-predictors and finding common major functional terms for their groups. Such groups of functions with molecular functions have the potential to provide a better explanation in CRISPR screens.

Overall our downstream analysis shows the reliability and sensibility of our models, which is directly associated with the prediction of the function of non-coding RNAs. The clustering of functions also highlighted the broader role of a few non-coding RNAs. For example, the noncoding RNA genes–ENSG00000259426, ENSG00000273063, ENSG00000224738, LINC00441, LINC01137, DLG1-AS1 were predicted to be associated with at least one of the members of the cluster of functions largely involved in cell cycle activity by our approach. Out of these 6 non-coding RNAs, DLG1-AS1 is reported to be involved in proliferation (Rui et al. 2018). The other two non-coding RNAs (LINC00441 (J. Zhou et al. 2018) and LINC01137 (Du et al. 2021) are reported to be involved in cancer development (Supporting File 5). Such inference about the role of non-coding RNAs in major processes could help biologists design experiments for validation.

Our approach of using the combination of epigenome and TFs as features and further clustering of functions to understand the role of coding and non-coding genes stands out of the crowd of gene-function prediction methods. We have created a resource for the biologists to corroborate their experiment results and utilize our predictions to design the experiments to understand the molecular and biological roles of non-coding and coding genes. The demonstration by our study advocates for the utilization of more epigenomic features for a better understanding of the functions of non-coding RNAs.

## Methods

The read-counts around promoters of genes were assimilated as described in Supplementary Methods. Notice here that in addition to TF and Histone modification ChIP-seq we also assimilated read-counts of a few input libraries without chromatin immunoprecipitation for different cell-type so that genome-wide bias can be captured and suppressed by ML models while learning the pattern for positive and negative sets of genes.

### Prediction method

For each gene-set in the ontology, we considered genes annotated in them as positive and randomly picked genes (not annotated in the same gene-set) as negatives, and gene function prediction is treated as a classification problem. Out of possible 50000 genes, if we expected the number of positive unknown genes belonging to a function is less than 100. Then if we were to pick 100 genes (as negatives) out of 50000 genes, there is less than 0.002 (100/500000) probability of having unknown positives in the negative set. For each gene-set, we choose an equal number of negatives to positives. We divided the positives and negatives into a training set (75%) and a test set (25%). The 5 different machine learning models for each gene-set are Random Forest, XGBoost, SVM (support vector machine), linear regression-based Lasso, and logistic regression.

Further, bootstrapping was done to calculate the standard deviation in the balanced accuracy by training the models for 5 iterations. We used various criteria to evaluate prediction on the test set, namely accuracy, balanced accuracy, F1-score and Mathew’s correlation coefficient (MCC), and error rate (Supporting File 1).

After evaluating the test set, we used the trained model to make predictions for all promoters (genes) in our list to find novel associations between function and genes. To have a stringent selection of novel/unknown gene-function associations predictions, we calculated the maximum true positive rate (TPR, named here as confidence score) for every function (gene-set).

#### Calculating confidence score for gene-sets

To have robust predictions, we calculated the confidence score of predictions for each function. For every function (gene-sets), each of their respective trained random forest models was used to predict probability scores (of belongingness to the gene-sets) for all 89747 promoters (genes). The confidence score of a gene-set is the threshold at which the probabilities of the genes yield the maximum (experimentally annotated) true genes against overall predicted genes. We have considered gene-sets having more than a 60% confidence score for our downstream analysis. The users can filter the gene-sets based on the confidence scores in our web server.

#### Inference about top regulators

We made inferences about top regulators by estimating feature importance while training random forest models. This approach has also been used by GENIE3, a top performer in genenetwork inference in the DREAM 5 challenge (Huynh-Thu et al. 2010; Marbach et al. 2012; Aibar et al. 2017). Here instead of gene expression of transcription factors, we are using binding affinity to promoters as feature scores, and we are predicting the belongingness of a gene to a class. Thus for every function, we chose the top 20 predictors with high feature importance calculated by the random forest-based approach.

### Method for Independent validations using PubMed abstract-mining

To gain confidence in novel predictions and compare our approach with other methods, we used PubMed abstract-based validation. Here, the ontology term and corresponding predicted gene term is used as input. In order to have a good match of the ontology term in a potentially relevant abstract, the ontology terms were processed to remove stop words (Supplementary Methods). The ‘*Bio.Entrez*’ package was used to search for the co-occurrence of the ontology term and its corresponding predicted gene term in abstracts of the research articles in the PubMed database. As a control to this approach, ontology function terms were paired randomly with gene terms and searched for their co-occurrence with the same parameters.

### Method for comparison of PubMed abstract mining result on predictions of different methods

A list of genes predicted from top-performing random forest models on 50 biological functions was used as input of other methods–Correlation AnalyzeR, DeepGO, GenetICA-Network, NetGO 2.0 (Miller and Bishop 2021; Kulmanov et al. 2018; Urzúa-Traslaviña et al. 2021; Yao et al. 2021). For the Correlation AnalyzeR method, the R library package ‘*correlationAnalyzeR*’ was used, ‘*analyzeSingleGenes*’ function from the package was used to predict the ontology-based labels for genes mentioned above predicted by our methods, ontology labels with the highest score were considered as the final predicted label. If the prediction was not available by the method for a gene, its label was left blank.

For DeepGO, GenetICA-Network, and NetGO 2.0 methods, their respective web servers were used to get the predictions on the considered list of genes by feeding the relevant protein sequence FASTA files as input; the top listed isoform was considered from the Uniport database (The UniProt Consortium 2017). The prediction label with non-generic terms with the highest score from either Biological Processes or Molecular Functions section was considered the final label. For non-coding genes and genes with less than a 50% score predictions, their labels were left blank.

PubMed abstract mining was run on all the predictions of different methods to get the cooccurrence of the predicted ontology term and input gene term using the ‘Bio.Entrez’ package as described above. Stop words (Supplementary Methods) were filtered out from the input terms to avoid matching generic terms.

### Method for clustering functions

To infer clusters of functions (gene-sets), we first estimated similarity scores among functions. The similarity score among the two functions was defined as the number of the same TFs and cofactor ChIP-seq (SRX id) profiles which appeared among the top-20 predictors for both. The similarity scores were inverted to get distances among functions to apply tSNE-based dimension reduction. For this purpose, R package ‘*Rtsne*’ was used with the option ‘is_distance’ equal to TRUE. After low dimensional embedding, DBSCAN was used to find clusters of functions using the 2D embedding coordinates provided by ‘*Rtnse*’ (Ester et al. 1996; Van der Maaten and Hinton 2008).

### Methodology for transcription factors synergy and pleiotropy analysis

The occurrence of TFs and cofactors as top 20 predictors for same and different cell-types across those biological functions with more than 60% confidence score was counted. Similarly, with the given set of TFs used as features. A TF pair list was constructed, and the occurrence of each TF pair among the top 20 predictors across the same biological functions was counted.

## Availability of data sources and code

Profiles of ChIP-seq of transcription factors, histone marks, and DNase-seq were downloaded from the ChipAtlas database (https://chip-atlas.org/) in bedGraph format, which can be processed by extension of the DFilter tool. The CAGE-tags profiles were downloaded from the FANTOM database (https://fantom.gsc.riken.jp/data/).

Peak counts of the epigenome profiles can be obtained using DFilter at https://reggenlab.github.io/DFilter/.

### Key Points

1. Good predictability for many functions (gene-set) using epigenome features shared between coding and non-coding RNA genes.
2. Unbiased validation and comparison with other methods for association predictions between gene and function.
3. Clustering functions based on shared predictors reveal their category in terms of major processes and corresponding top regulators.
4. Robust prediction and comprehensive analysis of combinatorics of regulators and clusters of function provide reliable insight into the role of non-coding RNAs.

## Supplementary Methods

### Feature construction

We used the read-counts of epigenome and transcriptome profiling assays as features. For this purpose, we counted the number of DNA fragments lying within 1 kbp of gene transcription start sites (TSS). We calculated the number of reads around TSS using ChIP-Seq (TF, Histone modifications) and DNAse-Seq profiles from the ChIP-Atlas database[1] and CAGE-tags from the FANTOM5 database. We also calculated the read-counts of a few input libraries (with ChIP) around the TSS of genes using an extension of tool DFilter[2]. The purpose of using the readcounts of the input library was to reduce the effect of artifacts and biases originating from the assays so that the predictability of functions can be improved. We used the TSS of non-coding genes from gencode (V30) and RefSeq gene transcripts[3,4]. For each gene, we allowed multiple transcripts as long as their TSS were at least 500 bp apart from each other. In total, we performed our analysis using 89747 promoter regions.

#### Balanced accuracy

It is a metric used to judge the predictive power of a binary classifier. It is often used when there is imbalance in the number of positive and negative (imbalanced classes). Balanced accuracy is defined as arithmetic mean of sensitivity and specificity:

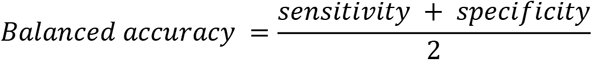

#### PubMed abstract mining stop words

These following stop-words were used to process the function terms to remove the generic terms. The following stop words were removed:

> Stop words 1: ‘cell’,‘of’,‘small’,‘in’,‘is’,‘he’, ‘to’, ‘from’, ‘by’, ‘on’, ‘the’, ‘or’, ‘like’, ‘layer’, ‘ii’, ‘groups’, ‘into’, ‘type’.
>
> Stop words 2: ‘binding’, ‘protein’, ‘factor’, ‘activity’, ‘regulation’, ‘group’, ‘chemical’, ‘sensory’, ‘other’, ‘process’, ‘species’, ‘positive’, ‘compound’, ‘cellular’, ‘particle’, ‘organism’, ‘involved’, ‘movement’, ‘interaction’, ‘environment’, ‘pathway’, ‘signaling’, ‘coupled’, ‘mrna’, ‘response’, ‘negative’,‘modified’, ‘response’, ‘left’, ‘right’, ‘formation’, ‘nucleotide’, ‘receptor’, ‘gene’, ‘complex’,‘dependent’, ‘maintenance’, ‘process’, ‘acid’.

### Method for comparison of PubMed abstract mining result on predictions of different methods

The following stop-words were removed to avoid matching with generic terms:

> Stop words 1: ‘cell’,‘of’,‘small’,‘in’,‘is’,‘he’, ‘to’, ‘from’, ‘by’, ‘on’, ‘the’, ‘or’, ‘like’, ‘layer’, ‘ii’, ‘groups’, ‘into’, ‘type’, ‘containing’, ‘protein’, ‘receptor’, ‘organ’.
>
> Stop words 2: = ‘binding’, ‘factor’, ‘activity’, ‘regulation’, ‘group’, ‘chemical’, ‘sensory’, ‘other’, ‘process’, ‘species’, ‘positive’, ‘compound’, ‘cellular’, ‘particle’, ‘organism’, ‘involved’, ‘movement’, ‘interaction’, ‘environment’, ‘pathway’, ‘signaling’, ‘coupled’, ‘mrna’, ‘response’, ‘negative’, ‘modified’, ‘response’, ‘left’, ‘right’, ‘formation’, ‘nucleotide’, ‘gene’, ‘complex’, ‘dependent’, ‘maintenance’, ‘process’, ‘acid’.

#### Method for validation using Mouse and human disease gene-sets

In order to further consolidate some of our predictions of novel associations between function and genes, we used information downloaded from the mouse disease-phenotype gene-sets available at the MGI database[5]. First, we found matching function terms of our ontology to which novel predictions are made, to mouse disease-phenotype terms, further, the intersection of predicted gene terms was done with the annotated mouse genes. If there is an intersection between function term and gene term with mouse disease and gene terms, then we call the prediction validated.

Similarly, we downloaded the gene-sets belonging to human diseases from the DisGeneNet database [6] and carried out the same method to validate the predictions.

### Supplementary Results

**Supplementary Figure 1:**
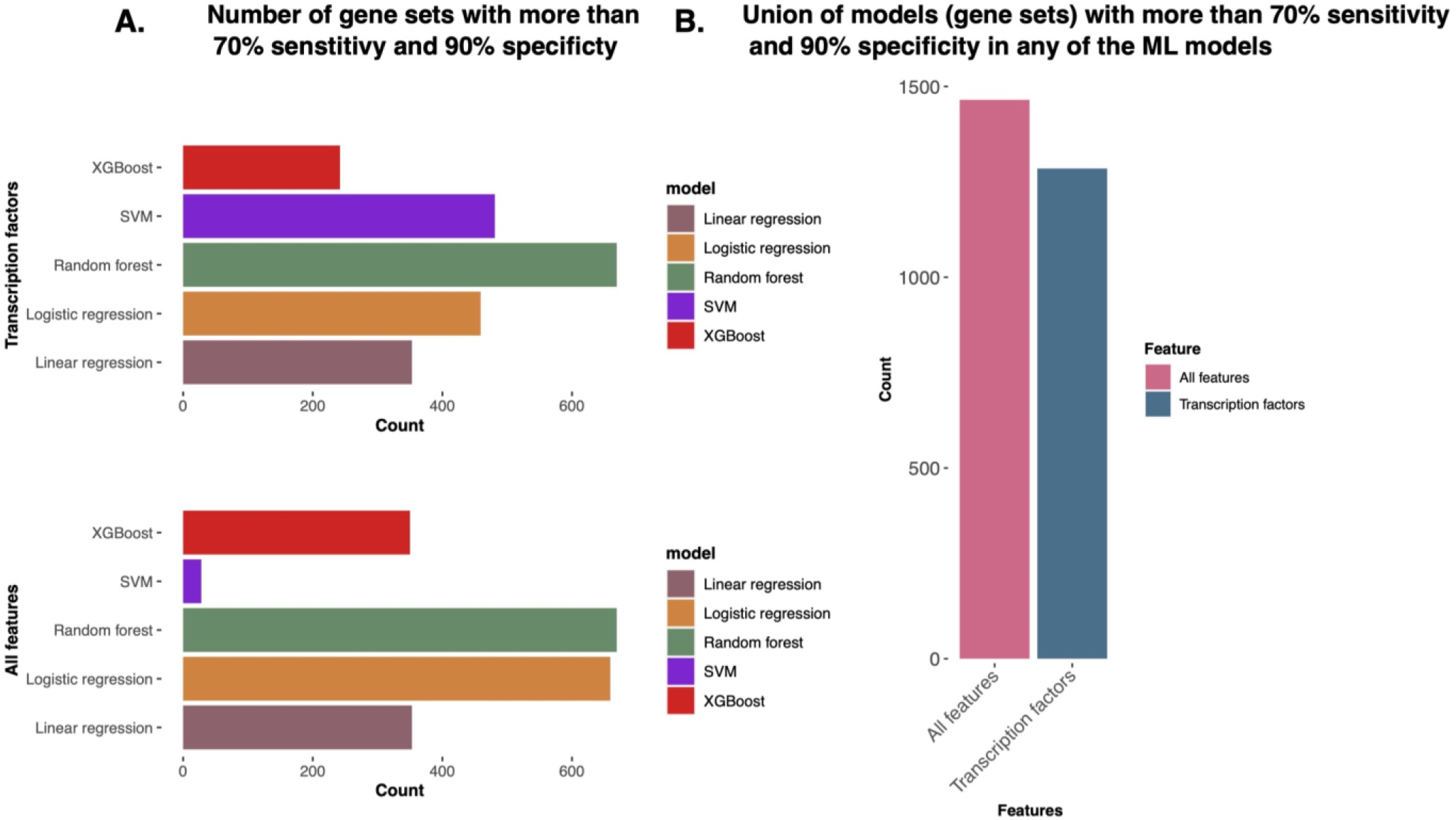
The Number of function (gene-sets) with satisfactory prediction (specificity 90%, sensitivity 70%) A) The number of gene-sets with satisfactory prediction by different machine learning (ML) methods. B) The number of union set of genes with satisfactory prediction by any of the 5 ML methods.

**Supplementary Figure 2:**
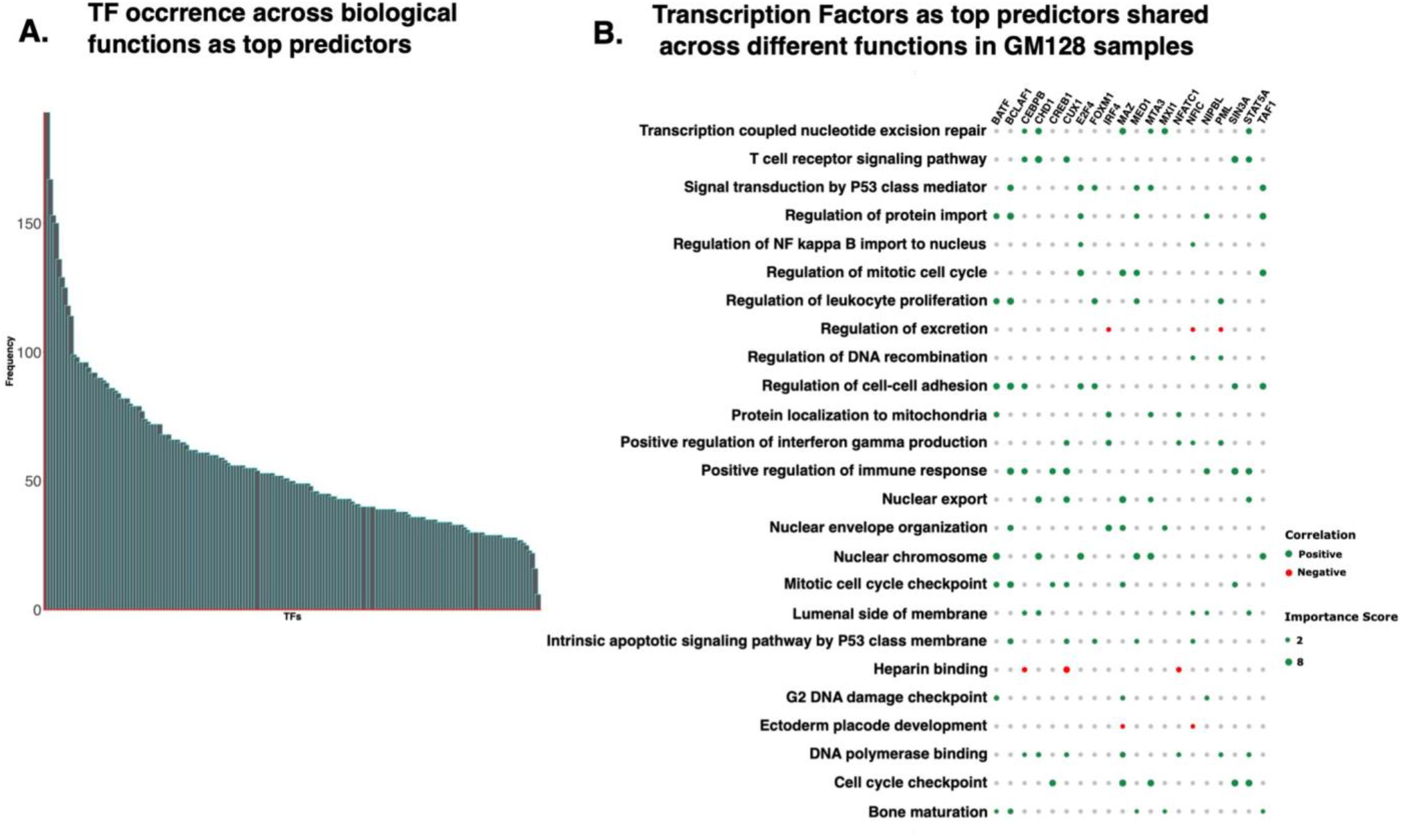
Predictive-pleiotropy of transcription factors (TFs) A) The number of functions where a TF appeared among top-predictor B) Dot-plot shows the feature importance score for TF-ChIP-seq profiles from GM12878 cells. The dot size shows the feature importance score and color shows the directionality according to Spearman correlation. Only those functions are shown for which at least 2 TF-ChIP-seq profiles from GM12878 cells were among the top 20 predictors.

**Supplementary Table 1.** Intersection of predicted genes with human disease-gene sets.

| Sl. No. | Disease term | Ontology term | Predicted Genes |
| --- | --- | --- | --- |
| 1 | leukocyte disorders | leukocyte mediated immunity | ITGB2 |
| 2 | male sterility due chromosome deletions | male gamete generation | RBMY1A1 |
| 3 | peripheral nervous system diseases | nervous | AR |
| 4 | organic mental disorders substance induced | organic catabolic | F12 |
| 5 | cardiac hypertrophy | cardiac muscle apoptotic | NPR 1.00 |
| 6 | metabolic myopathy | nucleoside monophosphate metabolic | ACADVL |
| 7 | global developmental delay | developmental maturation | NTNG2 |
| 8 | brain diseases metabolic inherited | dna metabolic | NDUFAF2 |
| 9 | disorder eye | eye morphogenesis | CDH3 |
| 10 | chromosomal instability | chromosomal region | KIF11 |
| 11 | chromosome 16p112 deletion syndrome kb | nuclear chromosome | SH2B1 |
| 12 | platelet abnormalities eosinophilia immune mediated inflammatory disease | activation immune | ARPC1B |
| 13 | acute myeloid leukemia m1 | myeloid leukocyte mediated immunity | CAPG |
| 14 | acute coronary syndrome | acute inflammatory | PON1 |
| 15 | metabolic bone disorder | steroid metabolic | C2 |
| 16 | neural tube defects | tube size | PYY |
| 17 | malignant lymphoma lymphocytic intermediate differentiation diffuse | lymphocyte differentiation | PIK3CD |
| 18 | B cells expansion nfkb anergy | B cells | CARD11 |
| 19 | lymphoma extranodal nk t cells | T cells | JAK3 |
| 20 | hodgkin lymphoma lymphocyte depletion | lymphocyte activation | TNF |
| 21 | cardiac arrhythmia | cardiac muscle tissue | KCNJ2 |
| 22 | abdominal obesity metabolic syndrome | organic hydroxy metabolic | MTTP |
| 23 | organic mental disorders substance induced | organic hydroxy metabolic | MSRA |

**Supplementary Table 2.**
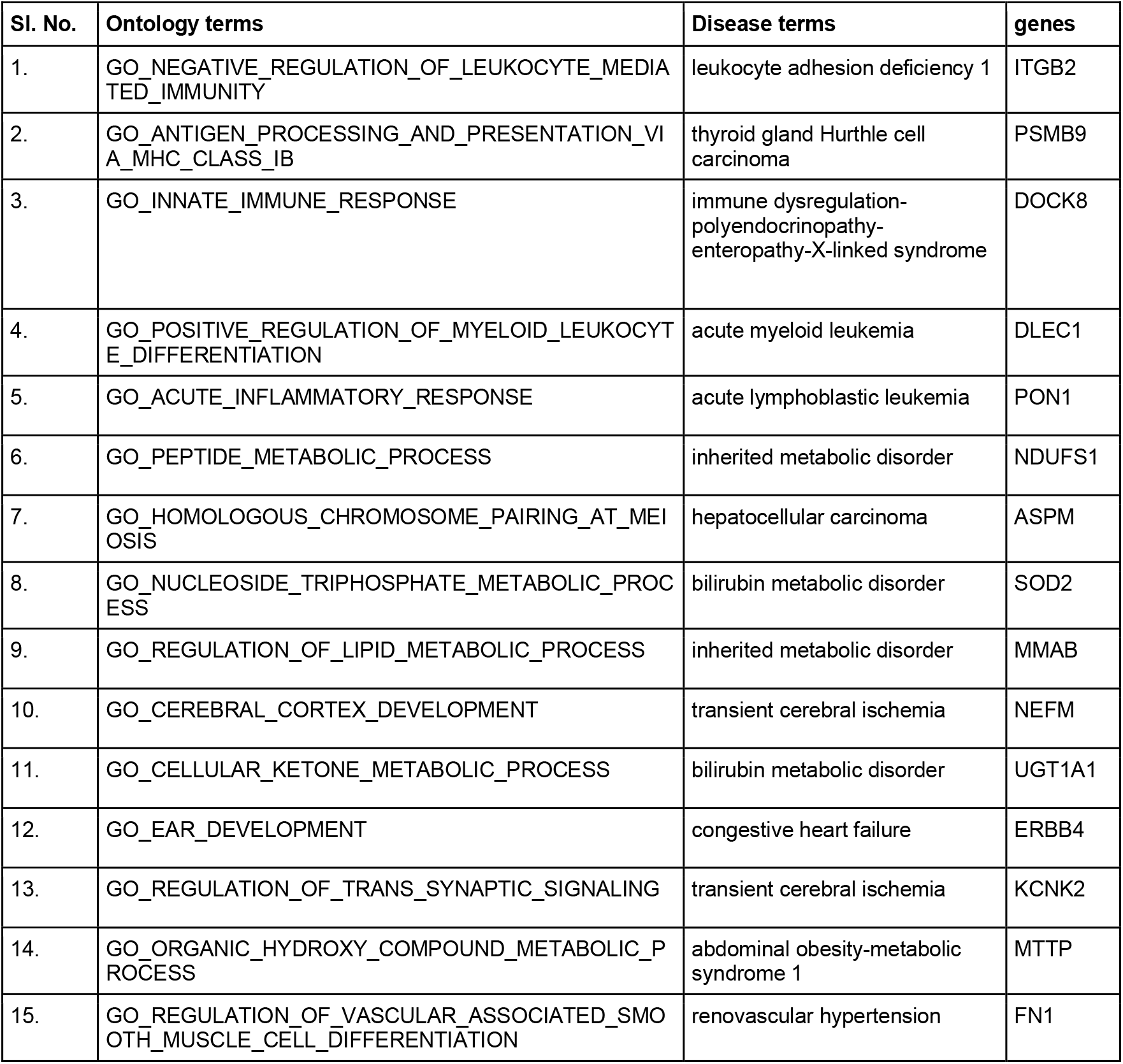
Intersection of predicted genes with mice disease-gene sets.

**Supplementary Table 3.** List of lncRNA genes with predicted function with literature evidence

| Ontology | Predicted non-coding gene | Literature evidence |
| --- | --- | --- |
| GO_MIDBRAIN_DEVELOPMENT | AC145422.1 | [7] |
| GO_CHROMATIN_ASSEMBLY_OR_DISASSEMBLY | ENSG00000260830 | [8] |
| GO_RNA_POLYMERASE_II_ACTIVATING_TRANSCRIPTION_FACTOR_BINDING | AC079305.10<br>(AC079305.1) | [9] |
| RNA_POLYMERASE_II_ACTIVATING_TRANSCRIPTION_FACTOR_BINDING | ENSG00000222043 | [10] |
| GO_CIRCADIAN_RHYTHM | AC024243.1 | [11] |
| GO_REGULATION_OF_CELL_ACTIVATION | AC007384.1 | [12] |
| GO_COVALENT_CHROMATIN_MODIFICATION | AC087623.3 | [13] |

## References

Aibar, Sara, Carmen Bravo González-Blas, Thomas Moerman, Vân Anh Huynh-Thu, Hana Imrichova, Gert Hulselmans, Florian Rambow, et al. 2017. “SCENIC: Single-Cell Regulatory Network Inference and Clustering.” Nature Methods 14 (11): 1083–86.

Alimov, Irina, Suchithra Menon, Nadire Cochran, Rob Maher, Qiong Wang, John Alford, John B. Concannon, et al. 2019. “Bile Acid Analogues Are Activators of Pyrin Inflammasome.” The Journal of Biological Chemistry 294 (10): 3359–66.

Bakhmet, Evgeny I., and Alexey N. Tomilin. 2021. “Key Features of the POU Transcription Factor Oct4 from an Evolutionary Perspective.” Cellular and Molecular Life Sciences: CMLS 78 (23): 7339–53.

Blake, J. A., R. Baldarelli, J. A. Kadin, J. E. Richardson, C. L. Smith, and C. J. Bult. 2021. “Mouse Genome Database (MGD): Knowledgebase for Mouse-Human Comparative Biology.” Nucleic Acids Research 49 (D1). https://doi.org/10.1093/nar/gkaa1083.

Chen, Jingcheng, Yizhuo Wang, Cong Wang, Ji-Fan Hu, and Wei Li. 2020. “LncRNA Functions as a New Emerging Epigenetic Factor in Determining the Fate of Stem Cells.” Frontiers in Genetics 11 (March): 277.

Chen, Yu-Xiao, Jie Ding, Wei-Er Zhou, Xuan Zhang, Xiao-Tong Sun, Xi-Ying Wang, Chi Zhang, et al. 2021. “Identification and Functional Prediction of Long Non-Coding RNAs in Dilated Cardiomyopathy by Bioinformatics Analysis.” Frontiers in Genetics 12 (April): 648111.

Chesmore, Kevin N., Jacquelaine Bartlett, Chao Cheng, and Scott M. Williams. 2016. “Complex Patterns of Association between Pleiotropy and Transcription Factor Evolution.” Genome Biology and Evolution 8 (10): 3159.

Donato, Luigi, Concetta Scimone, Simona Alibrandi, Carmela Rinaldi, Antonina Sidoti, and Rosalia D’Angelo. 2020. “Transcriptome Analyses of lncRNAs in A2E-Stressed Retinal Epithelial Cells Unveil Advanced Links between Metabolic Impairments Related to Oxidative Stress and Retinitis Pigmentosa.” Antioxidants (Basel, Switzerland) 9 (4). https://doi.org/10.3390/antiox9040318.

Du, Yong, Haiyan Yang, Yue Li, Wenli Guo, Yufeng Zhang, Haitao Shen, Lingxiao Xing, Yuehong Li, Wenxin Wu, and Xianghong Zhang. 2021. “Long Non-Coding RNA LINC01137 Contributes to Oral Squamous Cell Carcinoma Development and Is Negatively Regulated by miR-22-3p.” Cellular Oncology 44 (3): 595–609.

Elaine Hardman, W., Donald A. Primerano, Mary T. Legenza, James Morgan, Jun Fan, and James Denvir. 2019. “mRNA Expression Data in Breast Cancers before and after Consumption of Walnut by Women.” Data in Brief 25 (August): 104050.

Ester, M., H. P. Kriegel, J. Sander, and X. Xu. 1996. “A Density-Based Algorithm for Discovering Clusters in Large Spatial Databases with Noise.” KDD: Proceedings / International Conference on Knowledge Discovery & Data Mining. International Conference on Knowledge Discovery & Data Mining. https://www.aaai.org/Papers/KDD/1996/KDD96-037.pdf?source=post_page.

Hendricks, Kristin B., Frances Shanahan, and Emma Lees. 2004. “Role for BRG1 in Cell Cycle Control and Tumor Suppression.” Molecular and Cellular Biology 24 (1): 362–76.

Huynh-Thu, Vân Anh, Alexandre Irrthum, Louis Wehenkel, and Pierre Geurts. 2010. “Inferring Regulatory Networks from Expression Data Using Tree-Based Methods.” PloS One 5 (9). https://doi.org/10.1371/journal.pone.0012776.

Hyle, Judith, Yang Zhang, Shaela Wright, Beisi Xu, Ying Shao, John Easton, Liqing Tian, Ruopeng Feng, Peng Xu, and Chunliang Li. 2019. “Acute Depletion of CTCF Directly Affects MYC Regulation through Loss of Enhancer–promoter Looping.” Nucleic Acids Research 47 (13): 6699–6713.

Jeng, Edwin E., Varun Bhadkamkar, Nnejiuwa U. Ibe, Haley Gause, Lihua Jiang, Joanne Chan, Ruiqi Jian, et al. 2019. “Systematic Identification of Host Cell Regulators of Legionella Pneumophila Pathogenesis Using a Genome-Wide CRISPR Screen.” Cell Host & Microbe 26 (4): 551–63.e6.

Kenny, Colin, Elaine O’Meara, Mevlüt Ulaş, Karsten Hokamp, and Maureen J. O’Sullivan. 2021. “Global Chromatin Changes Resulting from Single-Gene Inactivation—The Role of SMARCB1 in Malignant Rhabdoid Tumor.” Cancers. https://doi.org/10.3390/cancers13112561.

Kevin C. Wang, Howard Y. Chang. 2011. “Molecular Mechanisms of Long Noncoding RNAs.” Molecular Cell 43 (6): 904.

Kim, Somi, Nam-Kyung Yu, and Bong-Kiun Kaang. 2015. “CTCF as a Multifunctional Protein in Genome Regulation and Gene Expression.” Experimental & Molecular Medicine 47 (6): e166–e166.

Kulmanov, Maxat, and Robert Hoehndorf. 2019. “DeepGOPlus: Improved Protein Function Prediction from Sequence.” Bioinformatics 36 (2): 422–29.

Kulmanov, Maxat, Mohammed Asif Khan, Robert Hoehndorf, and Jonathan Wren. 2018. “DeepGO: Predicting Protein Functions from Sequence and Interactions Using a Deep Ontology-Aware Classifier.” Bioinformatics 34 (4): 660–68.

Kumar, Vibhor, Masafumi Muratani, Nirmala Arul Rayan, Petra Kraus, Thomas Lufkin, Huck Hui Ng, and Shyam Prabhakar. 2013. “Uniform, Optimal Signal Processing of Mapped Deep-Sequencing Data.” Nature Biotechnology 31 (7): 615–22.

Liberzon, Arthur, Aravind Subramanian, Reid Pinchback, Helga Thorvaldsdóttir, Pablo Tamayo, and Jill P. Mesirov. 2011. “Molecular Signatures Database (MSigDB) 3.0.” Bioinformatics 27 (12): 1739–40.

Li, Bing, Michael Carey, and Jerry L. Workman. 2007. “The Role of Chromatin during Transcription.” Cell. https://doi.org/10.1016/j.cell.2007.01.015.

Liu, Guojun, Zihao Chen, Irina G. Danilova, Mikhail A. Bolkov, Irina A. Tuzankina, and Guoqing Liu. 2018. “Identification of miR-200c and miR141-Mediated lncRNA-mRNA Crosstalks in Muscle-Invasive Bladder Cancer Subtypes.” Frontiers in Genetics 0. https://doi.org/10.3389/fgene.2018.00422.

Li, Zhongyang, Shang Cai, Huijun Li, Jincheng Gu, Ye Tian, Jianping Cao, Dong Yu, and Zaixiang Tang. 2021. “Developing a lncRNA Signature to Predict the Radiotherapy Response of Lower-Grade Gliomas Using Co-Expression and ceRNA Network Analysis.” Frontiers in Oncology 11 (March): 622880.

Marbach, Daniel, James C. Costello, Robert Küffner, Nicole M. Vega, Robert J. Prill, Diogo M. Camacho, Kyle R. Allison, et al. 2012. “Wisdom of Crowds for Robust Gene Network Inference.” Nature Methods 9 (8): 796–804.

Meurer, Logan, Leonard Ferdman, Beau Belcher, and Troy Camarata. 2021. “The SIX Family of Transcription Factors: Common Themes Integrating Developmental and Cancer Biology.” Frontiers in Cell and Developmental Biology 9 (August): 707854.

Miller, Henry E., and Alexander J. R. Bishop. 2021. “Correlation AnalyzeR: Functional Predictions from Gene Co-Expression Correlations.” BMC Bioinformatics 22 (1): 206.

Pech, Matthew F., Linda E. Fong, Jacqueline E. Villalta, Leanne Jg Chan, Samir Kharbanda, Jonathon J. O’Brien, Fiona E. McAllister, Ari J. Firestone, Calvin H. Jan, and Jeffrey Settleman. 2019. “Systematic Identification of Cancer Cell Vulnerabilities to Natural Killer Cell-Mediated Immune Surveillance.” eLife 8 (August). https://doi.org/10.7554/eLife.47362.

Piñero, Janet, Àlex Bravo, Núria Queralt-Rosinach, Alba Gutiérrez-Sacristán, Jordi Deu-Pons, Emilio Centeno, Javier García-García, Ferran Sanz, and Laura I. Furlong. 2017. “DisGeNET: A Comprehensive Platform Integrating Information on Human Disease-Associated Genes and Variants.” Nucleic Acids Research 45 (D1): D833–39.

Raulerson, Chelsea K., Arthur Ko, John C. Kidd, Kevin W. Currin, Sarah M. Brotman, Maren E. Cannon, Ying Wu, et al. 2019. “Adipose Tissue Gene Expression Associations Reveal Hundreds of Candidate Genes for Cardiometabolic Traits.” American Journal of Human Genetics 105 (4): 773–87.

Rinn, John L., and Howard Y. Chang. 2012. “Genome Regulation by Long Noncoding RNAs,” June. https://doi.org/10.1146/annurev-biochem-051410-092902.

Rui, Xiaohui, Yun Xu, Yaqing Huang, Linjuan Ji, and Xiping Jiang. 2018. “lncRNA DLG1-AS1 Promotes Cell Proliferation by Competitively Binding with miR-107 and Up-Regulating ZHX1 Expression in Cervical Cancer.” Cellular Physiology and Biochemistry: International Journal of Experimental Cellular Physiology, Biochemistry, and Pharmacology 49 (5): 1792–1803.

Sage, Adam P., Kevin W. Ng, Erin A. Marshall, Greg L. Stewart, Brenda C. Minatel, Katey S. S. Enfield, Spencer D. Martin, Carolyn J. Brown, Ninan Abraham, and Wan L. Lam. 2020. “Assessment of Long Non-Coding RNA Expression Reveals Novel Mediators of the Lung Tumour Immune Response.” Scientific Reports 10 (1): 16945.

Sánchez-Jiménez, Carmen, Isabel Carrascoso, Juan Barrero, and José M. Izquierdo. 2013. “Identification of a Set of miRNAs Differentially Expressed in Transiently TIA-Depleted HeLa Cells by Genome-Wide Profiling.” BMC Molecular Biology 14 (February): 4.

Schinzel, Robert Thomas, Ryo Higuchi-Sanabria, Ophir Shalem, Erica Ann Moehle, Brant Michael Webster, Larry Joe, Raz Bar-Ziv, et al. 2019. “The Hyaluronidase, TMEM2, Promotes ER Homeostasis and Longevity Independent of the UPR.” Cell 179 (6): 1306–18.e18.

Sun, Xinghui, and Danny Wong. 2016. “Long Non-Coding RNA-Mediated Regulation of Glucose Homeostasis and Diabetes.” American Journal of Cardiovascular Disease 6 (2): 17–25.

Tak, Yu Gyoung, and Peggy J. Farnham. 2015. “Making Sense of GWAS: Using Epigenomics and Genome Engineering to Understand the Functional Relevance of SNPs in Non-Coding Regions of the Human Genome.” Epigenetics & Chromatin 8 (December): 57.

The UniProt Consortium. 2017. “UniProt: The Universal Protein Knowledgebase.” Nucleic Acids Research 45 (D1): D158–69.

Urzúa-Traslaviña, Carlos G., Vincent C. Leeuwenburgh, Arkajyoti Bhattacharya, Stefan Loipfinger, Marcel A. T. M. van Vugt, Elisabeth G. E. de Vries, and Rudolf S. N. Fehrmann. 2021. “Improving Gene Function Predictions Using Independent Transcriptional Components.” Nature Communications 12 (1): 1464.

Uygun, Sahra, Cheng Peng, Melissa D. Lehti-Shiu, Robert L. Last, and Shin-Han Shiu. 2016. “Utility and Limitations of Using Gene Expression Data to Identify Functional Associations.” PLoS Computational Biology 12 (12): e1005244.

Van der Maaten, Laurens, and Geoffrey Hinton. 2008. “Visualizing Data Using T-SNE.” Journal of Machine Learning Research: JMLR 9 (11). https://www.jmlr.org/papers/volume9/vandermaaten08a/vandermaaten08a.pdf?fbclid=IwA.

Venkatesh, Ishwariya, Vatsal Mehra, Zimei Wang, Matthew T. Simpson, Erik Eastwood, Advaita Chakraborty, Zac Beine, et al. 2021. “Co-Occupancy Identifies Transcription Factor Co-Operation for Axon Growth.” Nature Communications 12 (1): 2555.

Venters, B. J., and B. F. Pugh. 2013. “Genomic Organization of Human Transcription Initiation Complexes.” Nature 502 (7469). https://doi.org/10.1038/nature12535.

Wang, Zhi, Ben-Yang Liao, and Jianzhi Zhang. 2010. “Genomic Patterns of Pleiotropy and the Evolution of Complexity.” Proceedings of the National Academy of Sciences of the United States of America 107 (42): 18034–39.

Whittington, Camilla M., Denis O’Meally, Melanie K. Laird, Katherine Belov, Michael B. Thompson, and Bronwyn M. McAllan. 2018. “Transcriptomic Changes in the Pre-Implantation Uterus Highlight Histotrophic Nutrition of the Developing Marsupial Embryo.” Scientific Reports 8 (1): 2412.

Wu, Wei-Sheng, and Fu-Jou Lai. 2016. “Detecting Cooperativity between Transcription Factors Based on Functional Coherence and Similarity of Their Target Gene Sets.” PloS One 11 (9): e0162931.

Yang, Liu, Kun Huang, Xiangrao Li, Meng Du, Xiang Kang, Xi Luo, Lu Gao, et al. 2013. “Identification of Poly(ADP-Ribose) Polymerase-1 as a Cell Cycle Regulator through Modulating Sp1 Mediated Transcription in Human Hepatoma Cells.” PloS One 8 (12): e82872.

Yang, Peng, Xiao-Li Li, Jian-Ping Mei, Chee-Keong Kwoh, and See-Kiong Ng. 2012. “Positive-Unlabeled Learning for Disease Gene Identification.” Bioinformatics 28 (20): 2640–47.

Yang, Xinping, Jasmin Coulombe-Huntington, Shuli Kang, Gloria M. Sheynkman, Tong Hao, Aaron Richardson, Song Sun, et al. 2016. “Widespread Expansion of Protein Interaction Capabilities by Alternative Splicing.” Cell 164 (4): 805–17.

Yan, Jian, Yunjiang Qiu, André M. Ribeiro Dos Santos, Yimeng Yin, Yang E. Li, Nick Vinckier, Naoki Nariai, et al. 2021. “Systematic Analysis of Binding of Transcription Factors to Noncoding Variants.” Nature 591 (7848): 147–51.

Yao, Shuwei, Ronghui You, Shaojun Wang, Yi Xiong, Xiaodi Huang, and Shanfeng Zhu. 2021. “NetGO 2.0: Improving Large-Scale Protein Function Prediction with Massive Sequence, Text, Domain, Family and Network Information.” Nucleic Acids Research 49 (W1): W469–75.

Yilmaz, Atilgan, Mordecai Peretz, Aviram Aharony, Ido Sagi, and Nissim Benvenisty. 2018. “Defining Essential Genes for Human Pluripotent Stem Cells by CRISPR-Cas9 Screening in Haploid Cells.” Nature Cell Biology 20 (5): 610–19.

Zhang, Hanyu, Che-Lun Hung, Meiyuan Liu, Xiaoye Hu, and Yi-Yang Lin. 2019. “NCNet: Deep Learning Network Models for Predicting Function of Non-Coding DNA.” Frontiers in Genetics 0. https://doi.org/10.3389/fgene.2019.00432.

Zhang, Haoyue, Jessica Lam, Di Zhang, Yemin Lan, Marit W. Vermunt, Cheryl A. Keller, Belinda Giardine, Ross C. Hardison, and Gerd A. Blobel. 2021. “CTCF and Transcription Influence Chromatin Structure Re-Configuration after Mitosis.” Nature Communications 12 (1): 1–16.

Zhang, Xiaopei, Wei Wang, Weidong Zhu, Jie Dong, Yingying Cheng, Zujun Yin, and Fafu Shen. 2019. “Mechanisms and Functions of Long Non-Coding RNAs at Multiple Regulatory Levels.” International Journal of Molecular Sciences 20 (22): 5573.

Zhao, Yingwen, Jun Wang, Jian Chen, Xiangliang Zhang, Maozu Guo, and Guoxian Yu. 2020. “A Literature Review of Gene Function Prediction by Modeling Gene Ontology.” Frontiers in Genetics 0. https://doi.org/10.3389/fgene.2020.00400.

Zhou, Jianping, Jun Shi, Xingli Fu, Boneng Mao, Weimin Wang, Weiling Li, Gang Li, and Sujun Zhou. 2018. “Linc00441 Interacts with DNMT1 to Regulate RB1 Gene Methylation and Expression in Gastric Cancer.” Oncotarget 9 (101): 37471–79.

Zhou, Naihui, Yuxiang Jiang, Timothy R. Bergquist, Alexandra J. Lee, Balint Z. Kacsoh, Alex W. Crocker, Kimberley A. Lewis, et al. 2019. “The CAFA Challenge Reports Improved Protein Function Prediction and New Functional Annotations for Hundreds of Genes through Experimental Screens.” Genome Biology 20 (1): 244.

Zhu, Lihua J., Claude Gazin, Nathan D. Lawson, Hervé Pagès, Simon M. Lin, David S. Lapointe, and Michael R. Green. 2010. “ChIPpeakAnno: A Bioconductor Package to Annotate ChIP-Seq and ChIP-Chip Data.” BMC Bioinformatics 11 (May): 237.

Zhu, Ping, Jing Pan, Qian Qian Cai, Fan Zhang, Min Peng, Xing Li Fan, Hua Ji, Yi Wei Dong, Xing Zhong Wu, and Li Hui Wu. 2022. “MicroRNA Profile as Potential Molecular Signature for Attention Deficit Hyperactivity Disorder in Children.” Biomarkers: Biochemical Indicators of Exposure, Response, and Susceptibility to Chemicals, February, 1–10.

## Supplementary References

1. Oki S, Ohta T, Shioi G, et al. ChIP-Atlas: a data-mining suite powered by full integration of public ChIP-seq data. EMBO Rep. 2018; 19:

2. Kumar V, Muratani M, Rayan NA, et al. Uniform, optimal signal processing of mapped deepsequencing data. Nat. Biotechnol. 2013; 31:615–622

3. Frankish A, Diekhans M, Jungreis I, et al. GENCODE 2021. Nucleic Acids Res. 2021; 49:D916–D923

4. O’Leary NA, Wright MW, Brister JR, et al. Reference sequence (RefSeq) database at NCBI: current status, taxonomic expansion, and functional annotation. Nucleic Acids Res. 2016; 44:D733–45

5. Blake JA, Baldarelli R, Kadin JA, et al. Mouse Genome Database (MGD): Knowledgebase for mouse-human comparative biology. Nucleic Acids Res. 2021; 49:

6. Piñero J, Bravo À, Queralt-Rosinach N, et al. DisGeNET: a comprehensive platform integrating information on human disease-associated genes and variants. Nucleic Acids Res. 2017; 45:D833–D839

7. Saba LM, Hoffman PL, Homanics GE, et al. A long non-coding RNA (Lrap) modulates brain gene expression and levels of alcohol consumption in rats. Genes Brain Behav. 2021; 20:e12698

8. Li Z, Cai S, Li H, et al. Developing a lncRNA Signature to Predict the Radiotherapy Response of Lower-Grade Gliomas Using Co-expression and ceRNA Network Analysis. Front. Oncol. 2021; 11:622880

9. Lin W, Wang Y, Chen Y, et al. Role of Calcium Signaling Pathway-Related Gene Regulatory Networks in Ischemic Stroke Based on Multiple WGCNA and Single-Cell Analysis. Oxid. Med. Cell. Longev. 2021; 2021:8060477

10. Todoerti K, Ronchetti D, Favasuli V, et al. DIS3 mutations in multiple myeloma impact the transcriptional signature and clinical outcome. Haematologica 2021;

11. Wilsbacher LD, Sangoram AM, Antoch MP, et al. The mouse Clock locus: sequence and comparative analysis of 204 kb from mouse chromosome 5. Genome Res. 2000; 10:1928–1940

12. Huang Q-R, Pan X-B. Prognostic lncRNAs, miRNAs, and mRNAs Form a Competing Endogenous RNA Network in Colon Cancer. Front. Oncol. 2019; 9:712

13. Goldrich DY, LaBarge B, Chartrand S, et al. Identification of Somatic Structural Variants in Solid Tumors by Optical Genome Mapping. J Pers Med 2021; 11:

